# Genome-wide translocation events drive the evolution of *Candida africana*

**DOI:** 10.1101/2022.04.23.489250

**Authors:** Nnaemeka Emmanuel Nnadi, Domenico Giosa, Yocy Izam, Amaka Ubani, Anayochukwu Ngene, Grace Mebi Ayanbimpe, Ifeoma Bessie Enweani-Nwokelo, John Chinyere Aguiyi

## Abstract

*Candida africana* is emerging as an organism of interest. It is evolutionarily divergent from *Candida albicans* but has reduced virulence with a restricted ecological niche. This study aimed at comparing in silico the genome level to detect variations in the two species. Raw Illumina Hiseq data were downloaded from the European Nucleotide Archive (https://www.ebi.ac.uk/ena) with the accession number <u>SRR6669859</u> and assembled using shovill (v. 1.0.9) the resulting genome was mapped against the haploid reference *Candida albicans* SC5314_A22 strain using the D-GENIES webtool and contigs were reordered based on the reference, then gap-filled using GapFiller (v1-10), and annotated using MAKER-P. Synima and progressive Mauve were used to compare the annotated genomes of *Candida africana* and *Candida albicans* for synteny. OrthoVenn2 webserver was used for the identification and comparison of orthologous clusters. Microsynteny variations within the genomes were determined using the GEvo. The study revealed the presence of insertions, deletions, and hypervariable regions within the genome of *Candidia africana*, showing a high level of synteny with *Candida albicans*. The genome of *Candida africana* is 14.04Mbp with a BUSCO score of 99.66%. The two species form a total of 5146 orthologous protein clusters and shared a total of 5124 protein, *C. africana* has a unique cluster protein cluster while *C. albicans* have 18 unique Protein clusters. The genome of *C. africana* has lots of structural variations and the presence of gene losses and gains. These genetic variations possibly play a role in the reduced virulence potential observed in *C. africana*.

## Introduction

The epidemiology of *Candida* species causing infections is changing, with an increasing occurrence of non-albicans *Candida*. Despite this changing trend(1), *C. albicans* still remains the most dominant largely because of its genetic plasticity(2). *Candida africana* is the most evolutionary divergent lineage currently known within the *C. albicans* population(3). Ropars et al. (4)showed that *C. africana* is genetically different from *C. albicans*, having a low number of SNPs within the clade compared with *C. albicans* from other clades. Mixão and Gabaldón (5)presented that *C. africana* is however a product of evolution that has occurred in *C. albicans*. In their study,Ropars, Maufrais (4) showed that *C. africana* has 39 open reading frames (ORF) with premature stop codons (deleterious mutations) that were fixed in the clade. The ORFs contain genes encoding transcription factors required for fitness in systemic infection and proper regulation of morphogenesis, such as SFL1 (6) and ZCF29(7). The study proved that *C. africana* has much lower heterozygosity compared to other strains of C. *albicans*. This may be due to a combination of ancestral Loss of heterozygosity (LOH) and clonal reproduction with fixation of several deleterious alleles, affecting the overall fitness of these strains and leading to its niche restriction.

Genome plasticity is common in all eukaryotes(8). This can take the form of DNA insertions and deletions (indels), copy number variations (CNV), and loss of heterozygosity (LOH), which are frequently described during the evolution of organisms and disease states, such as cancer. No true meiosis has been observed in the most prevalent human fungal pathogen, *C. albicans*. Instead, it undergoes a parasexual process that involves random chromosome loss and rare Spo11-dependent chromosome recombination events(9). Despite its clonal lifestyle, *C. albicans* isolates exhibit extensive genomic diversity in the form of *de novo* base substitutions, indels, ploidy variation (haploid, diploid, and polyploid), karyotypic variation due to segmental and whole chromosome aneuploidies, and allele copy number variation including LOH(10). LOH events can occur over long ranges of the chromosome resulting in homozygous genes that could yield unexpected phenotypes such as attenuated virulence(11).

The impact and effects of LOH have been not understood in *C. africana*, a pathogen with different phenotypic and genotypic characteristics with reported inability to produces chlamy-dospores or assimilate N-acetylglucosamine. N-acetylglucosamine stimulates hyphal cell morphogenesis, virulence genes, and the genes needed to catabolize *N*-acetvigiucosamine in *C. albicans(12)*. *C. africana* has been shown to have a reduced formation of hyphae(13). The slow formation of hyphae has been suggested as a possible reason for reduced virulence when compared with *C. albicans* in a *Galleria mellonella* model(13). To date, there is no report of *C. africana* from the environment, hence its ecological niche is unknown but the report of a similar *Candida* belonging to the clade 13 [2] suggests that the environment may be the ecological niche of this yeast. This study aimed at comparing in silico the genomes of *Candida africana* and *Candida albicans* to identify variations in the genome of *C. africana* that could impact on the virulence of *C. africana*

## MATERIALS AND METHODS

### GENOME ASSEMBLY AND ANNOTATION

Raw Illumina Hiseq data were downloaded from the European Nucleotide Archive (https://www.ebi.ac.uk/ena) with the accession number SRR6669859. Adapter and low-quality reads were removed using Trimmomatic (v.0.39)(14). The remaining good-quality reads were analyzed with kat (v.2.4.2)(15) to investigate heterozygous ratio and then assembled with shovill (v.1.0.9)(16). Using a range of Kmer values. The homozygous genome was produced using redundans (v.0.14a)(17) on shovill’s contigs. The resulting genome was mapped against the haploid reference *Candida albicans* SC5314_A22 strain using the D-GENIES webtool (http://dgenies.toulouse.inra.fr), and contigs were reordered based on the reference. GapFiller (v1-10)(18) was used for the gap closure step, while the quality and completeness of the final genome were then evaluated using QUAST (19) and FGMP (20)

The assembled genome was annotated using MAKER-P (21)using mRNA-seq data for *Candida africana(22)*, Uniprot protein(23) for the *Candida* genus. Repeatmasker(24) was used to mask repeats, and gene prediction was made after training SNAP twice(25). To prepare the annotated genome for submission, wgs2ncbi(26)was used. Busco 3.1.0 (https://usegalaxy.eu) was used to inspect the quality of the final genome, using the following databases (ascomycota_odb9).

### COMPARATIVE GENOMICS

For comparisons of sequences at the genome level and to detect orthologs synteny between the two genomes, contigs of *Candida africana* were aligned with the reference sequences of the *Candida albicans* SC5314(vs22) using synima (27). Orthologues can be used to provide evidence for synteny, i.e. looking at the conservation of the ordering of loci on chromosomes between two individuals or species(27). To evaluate the assembly for completeness we used mummer (28). To detect the effects of gene re-arrangements, segmental duplication, gain or loss of gene within the genome of *Candida africana*, the genomes of the two organisms were visualized using progressivemauve (29). Microsynteny variations within the genomes were determined using the CoGe

#### Structural variation detection

To detect structural variations between the assembled *C. africana* genome and *Candida albicans*, the *C. africana* genome was aligned to the reference genome using nucmer(28), and the results were analyzed using and analyzed on the assemblytics(30).

#### Orthologous cluster determination

The annotated protein sets of *Candida africana* and the protein sequences of *Candida albicans* were analyzed using the OrthoVenn2(31) online web server (https://orthovenn2.bioinfotoolkits.net) for the identification and comparison of orthologous clusters. Briefly, to identify orthologous groups, OrthoVenn2 employs the OrthoMCL(32) clustering algorithm to annotate and compare ortholog groups, performing an all-against-all alignment using DIAMOND v0.9.24(33). The Gene Ontology (GO) terms for biological process, molecular function, and cellular component categories were assigned to the corresponding orthologous cluster by identifying similarity to sequences in the Uniprot (23) database. The e-value cutoff for all-to-all protein similarity comparisons was 0.05 and the inflation value for the generation of orthologous clusters using the Markov Cluster Algorithm was 1.5.

## RESULTS

A total of 20,612,494 raw reads were produced by the Illumina sequencing, and over 96.83% of those remained of good-quality for the downstream analysis. After the use of shovill, dgenies and redundans[17], we produced the homozygous genome assembly of *C. africana* consisting of 8 nuclear and a single mitochondrial chromosomes, with a total size of 14.04Mbp and a genome coverage of 176X. The genome annotation obtained using Maker-P[21] produced a total number of 3578 protein-coding genes, 94.9% of those assigned by FGMP (Table1).

**Table 1.** Fungal Highly conserved elements (FHCEs) report for Candida africana CBS11016 strain using FGMP

| **ASSEMBLY PARAMETER** | **ASSEMBLY STATISTICS** |
| --- | --- |
| **NUM OF UCES FOUND** | 28 out of 31 |
| **UCES COMPLETION ESTIMATION** | 90.3% |
| **TOTAL NUM OF PREDS ANALYZED** | 3304 |
| **NO. BEST FILTERED PREDS** | 455 |
| **NUM OF MARKERS COMPLETE** | 360 |
| **NUM OF MARKERS PARTIAL:** | 95 |
| **NUM OF ABERRANT PROTEINS** | 16 |
| **NUM OF FGMP COMPLETENESS MARKERS** | 563 |
| **ESTIMATE COMPLETENESS** | 94.9% |
| **LARGEST CONTIG** | 3150431 |
| **GC (%)** | 33.34 |
| **N50** | 2146907 |
| **NG50** | 2146907 |
| **N75** | 1609384 |
| **L50** | 3 |
| **LG50** | 3 |
| **L75** | 5 |
| **LG75** | 5 |

Key:

Proteins = 593 conserved fungal genes
DNA = 31 fungal highly conserved elements
%completeness = percent of 593 FCGs in the dataset

Orthologous genes are clusters of genes in different species that have evolved by vertical descent from a single ancestral gene (Fig.5). It results in 1619 protein clusters in *C. africana*, with 28 singleton proteins, meanwhile *C. albicans* has 6221 protein clusters with 3572 singletons. Among the clusters, *C. albicans* and *C. africana* shared 5127 clusters, with a single cluster unique to *C. africana* and 18 clusters unique to *C. albicans*.

**Figure 1:**
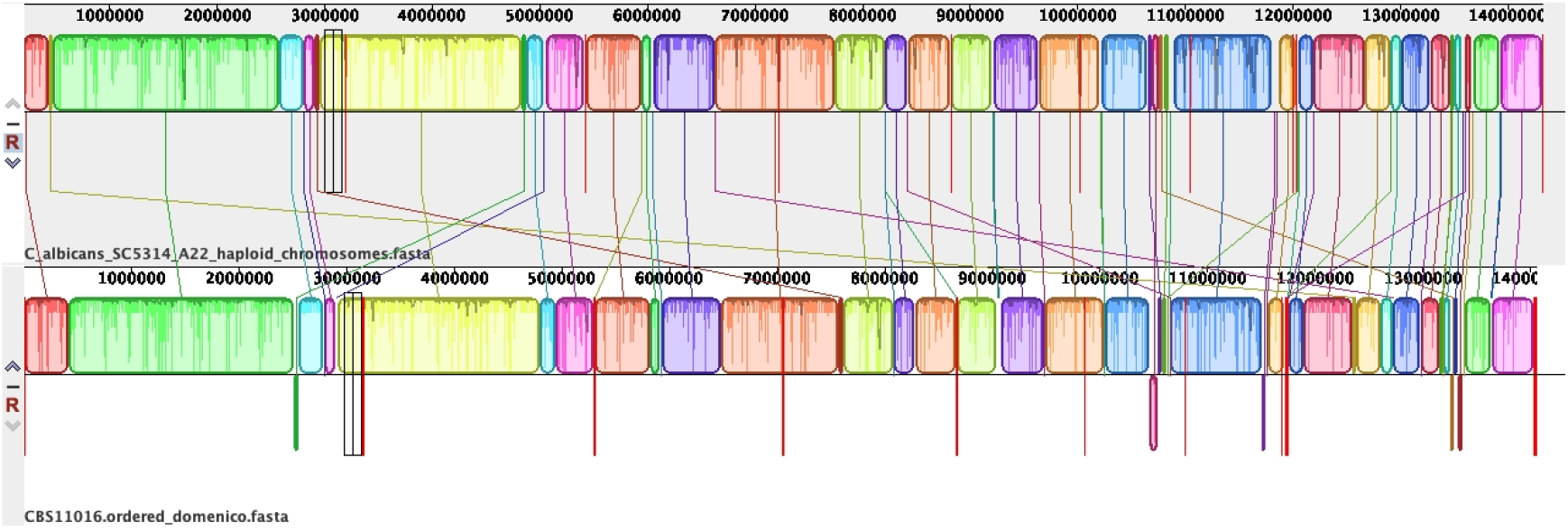
Structural organization of the genomes of *Candida africana* and *Candida albicans* SC5314. The color blocks (known as Locally Collinear Blocks) are conserved segments of sequences internally free from genome rearrangements.

**Figure 2:**
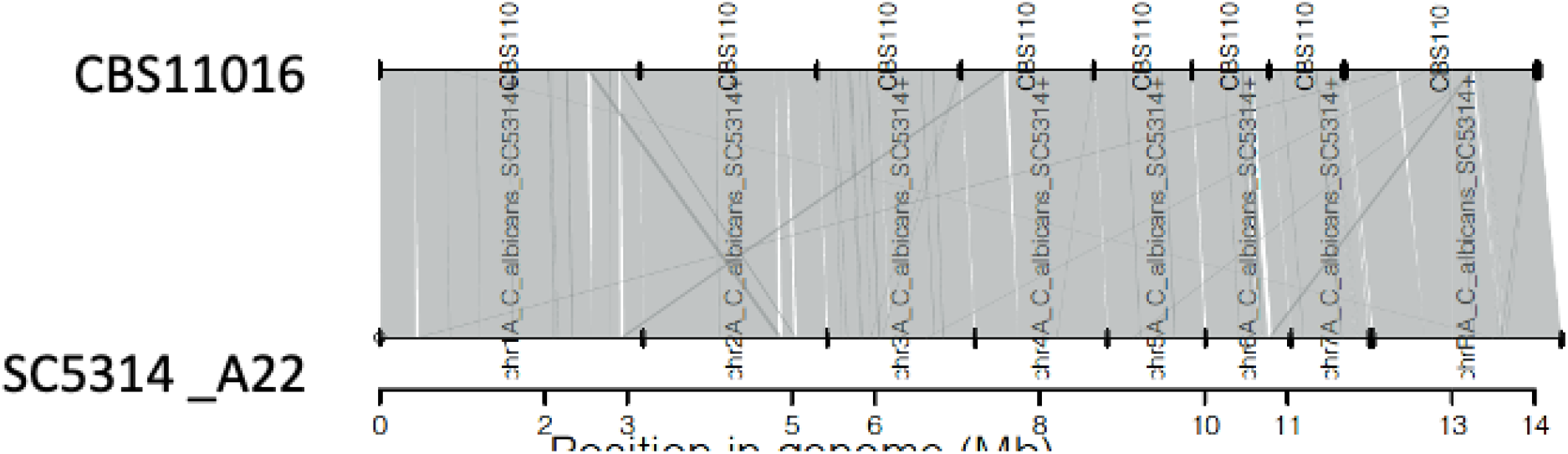
Synteny plot

**Figure 3:**
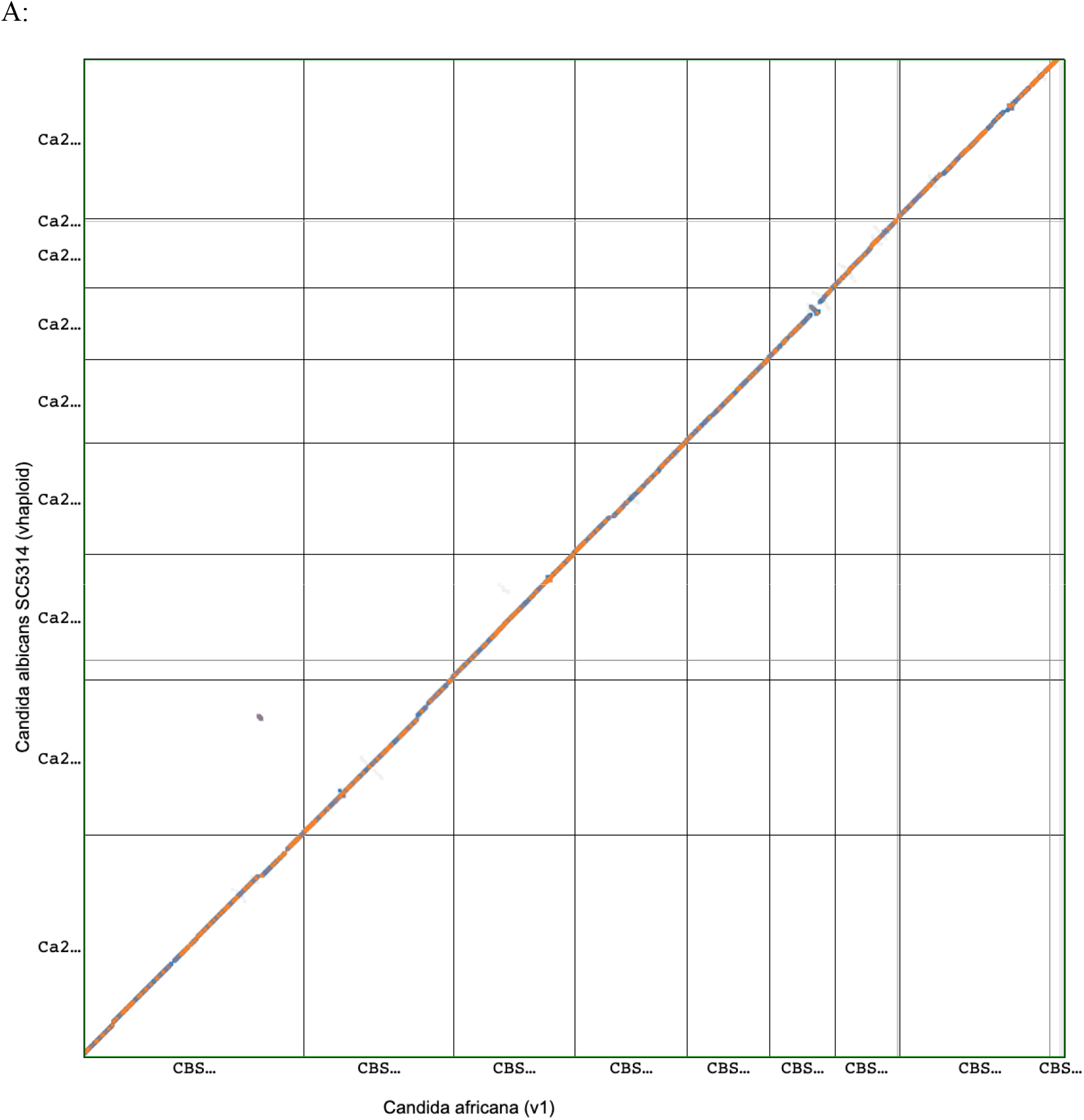
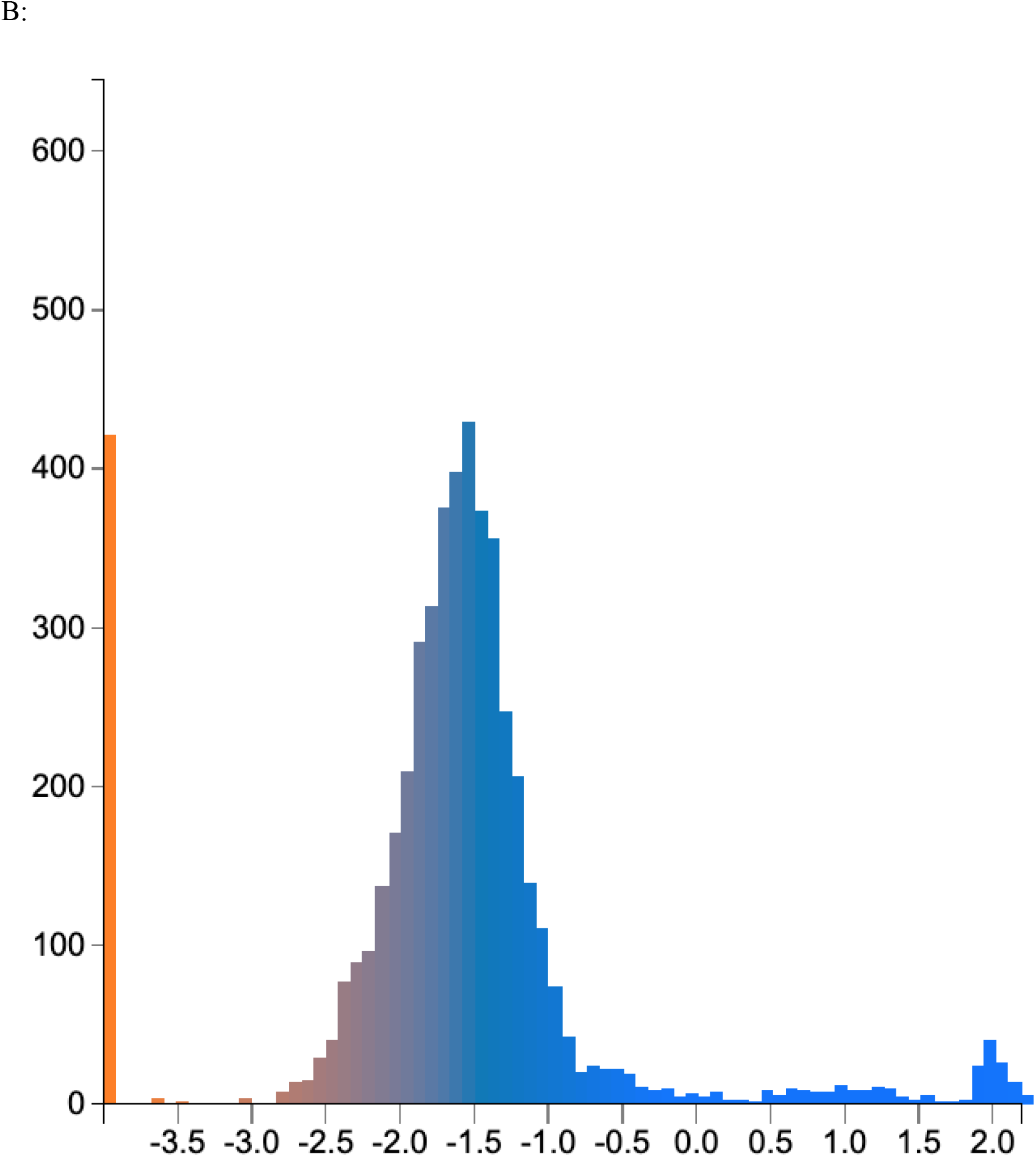
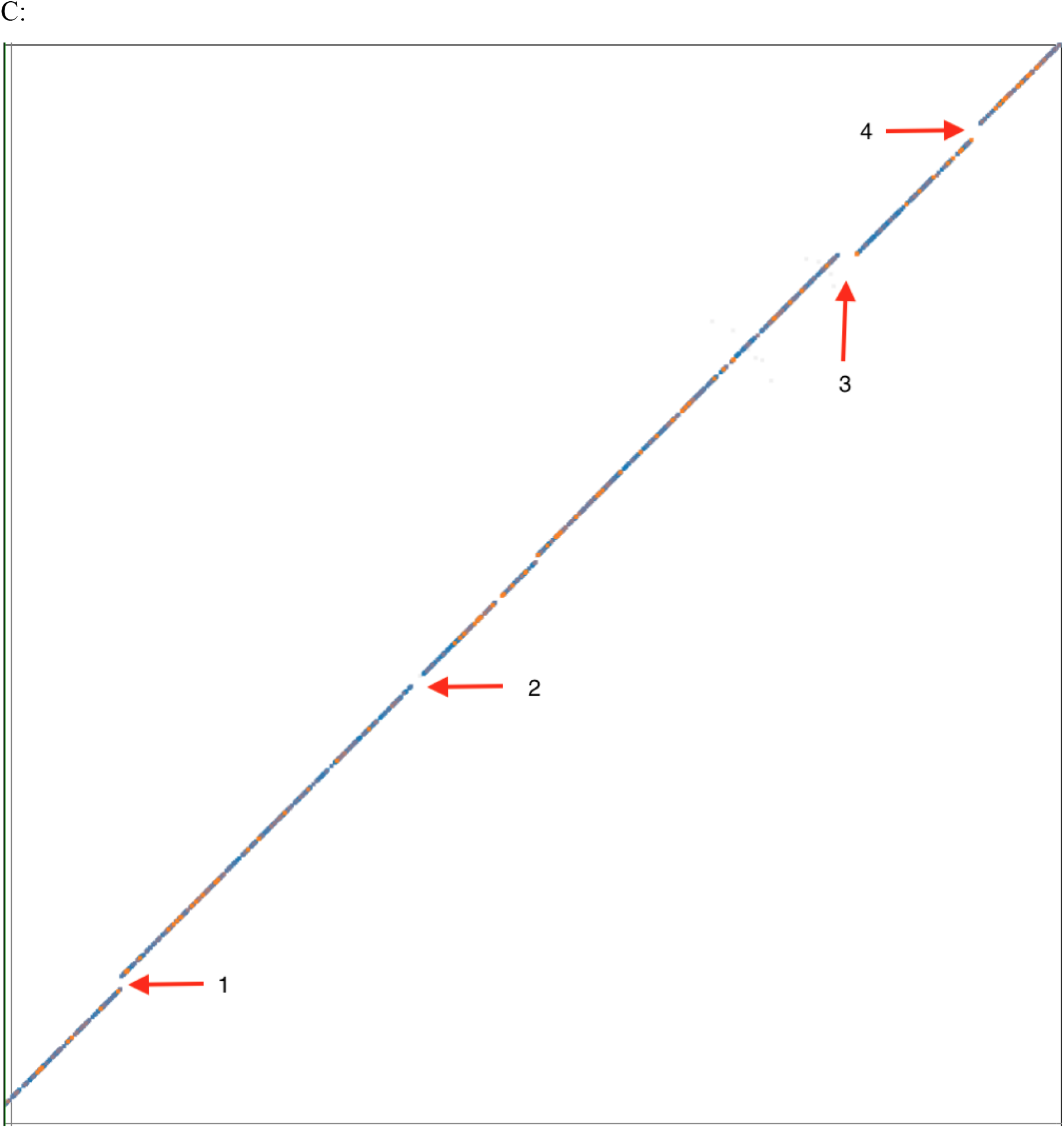
Syntenic dot plots and Ks plot between *Candida albicans* and *Candida africana*. Results may be regenerated: **A**. Whole-genome comparison between these two *Candida* species; *C. africana* on the x-axis and *C. albicans* on the y-axis. Horizontal and vertical grey lines separate chromosomes; colored dots represent syntenic gene pairs. Overall, the two genomes have very similar genome structures. **B**. Histogram of log10 transformed Ks values of syntenic gene pairs. Colors in the histogram are used to color syntenic gene pairs in the dot plots (**A and C).** The orange line represents gene-pairs with no synonymous mutations followed by a population of genes with 0.01 to 0.1 synonymous mutations per synonymous site. **C.** Syntenic dot plot of chromosome 1 of the two genomes. While the overall genomic structure is highly conserved, there are discontinuities in the syntenic regions representing regions of the genome with insertions, deletions, and hypervariable regions. The regions with an arrow and number are used for microsynteny analyses in Figure 4.

**Figure 4.1.**
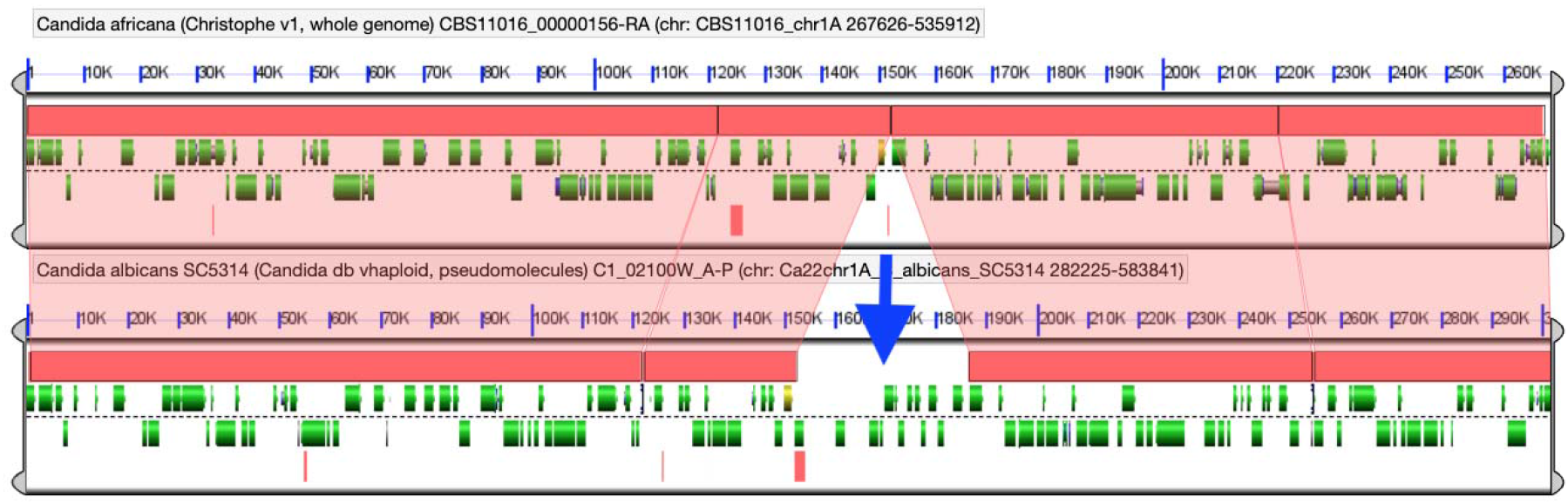
*C. albicans* region of around 30Kbp missing in *C. africana* This could be potentially the result of an insertion in *C. albicans* or a deletion in *C. africana*.

**Figure 4.2.**
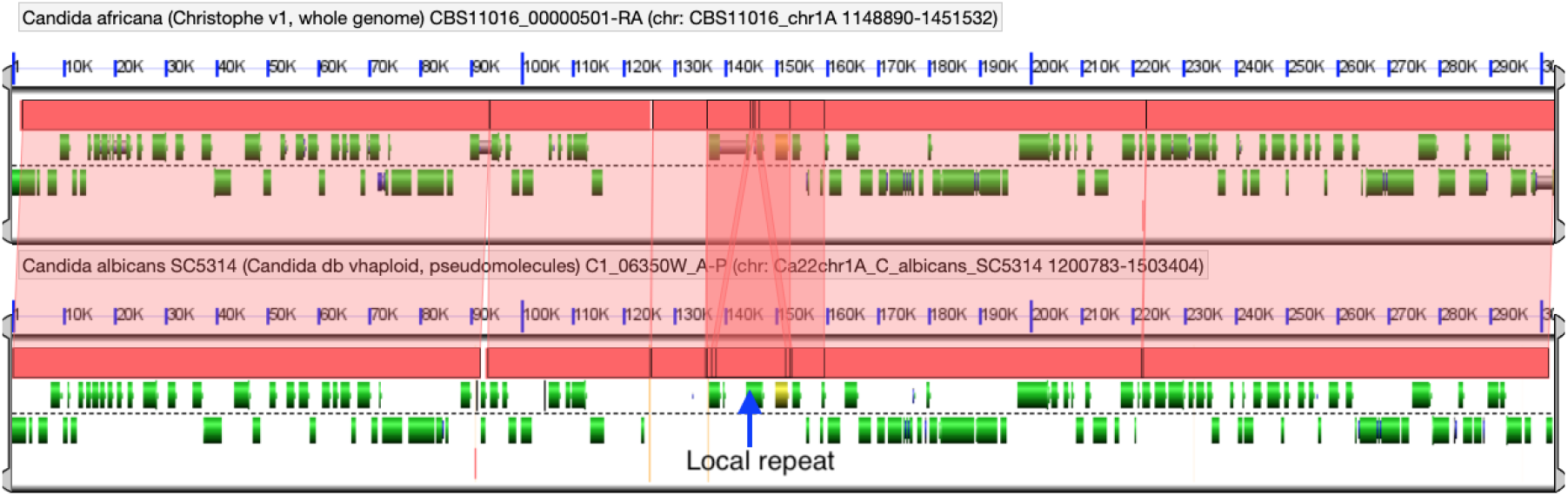
Syntenic discontinuity is caused by a local repeat region

**Figure 4.3.**
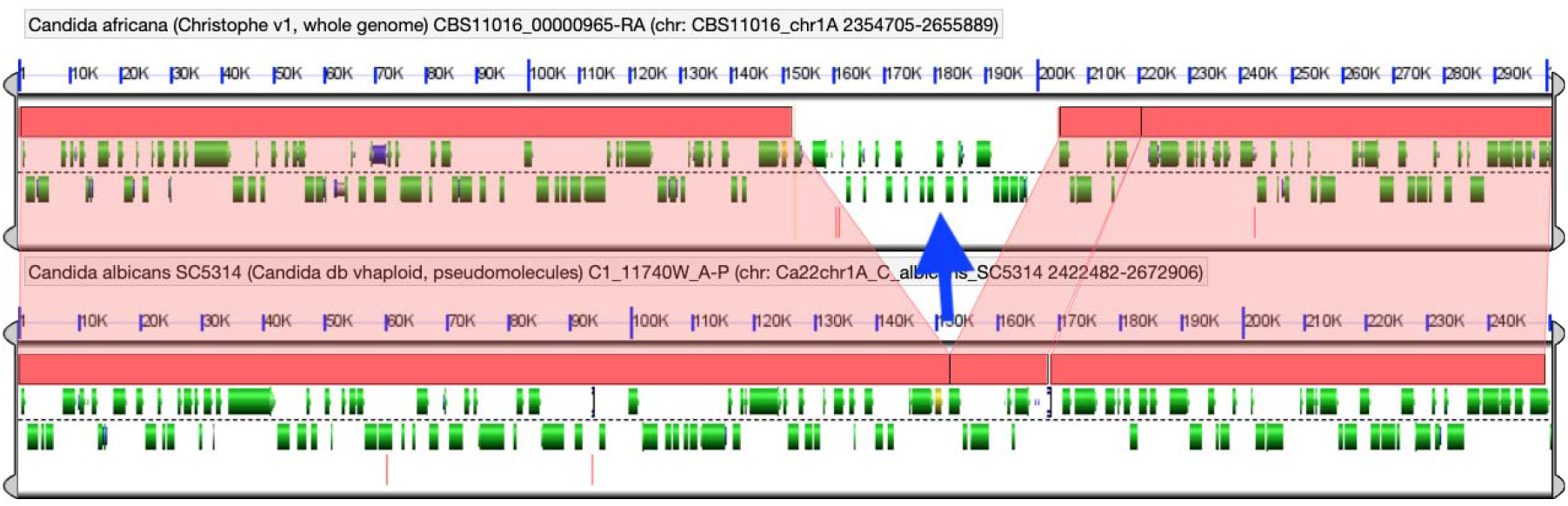
*C. africana* region of around 50kbp not present in *C. albicans* this may be the result of an insertion in *C. africana* or a deletion in *C. albicans*.

**Figure 4.4.**
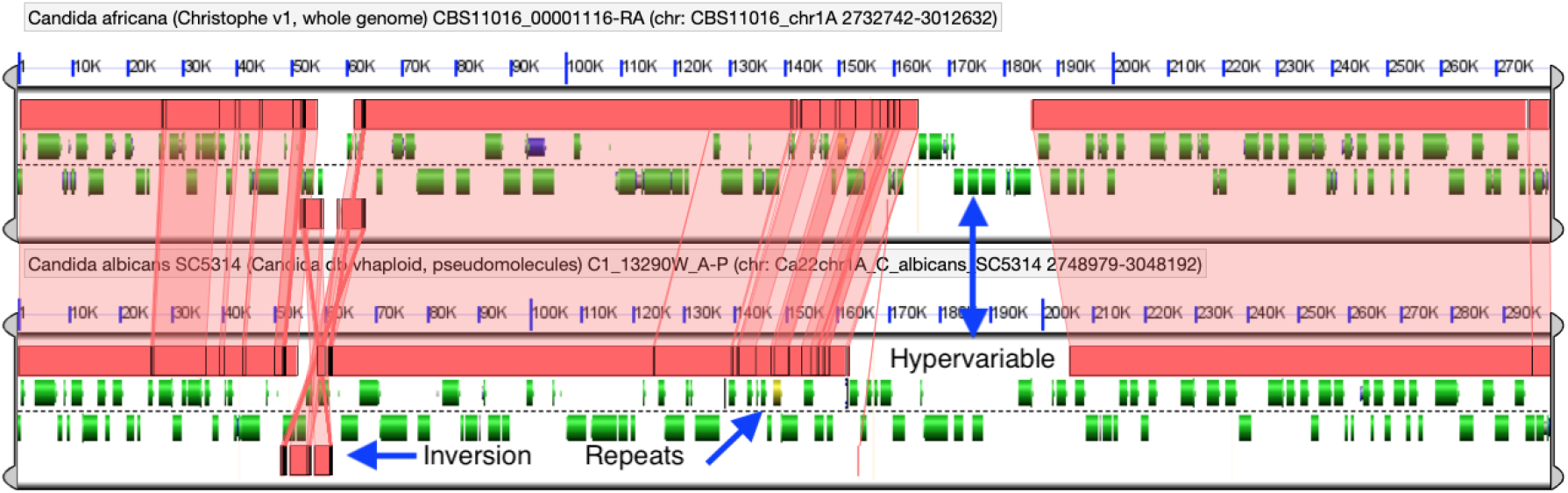
A complex set of genomic features with a hypervariable region in both genomes showing no mutual sequence similarity. In this genomic coordinates are showed also repetitive sequences near the hypervariable region, and a neighboring inversion

**Figure 5.**
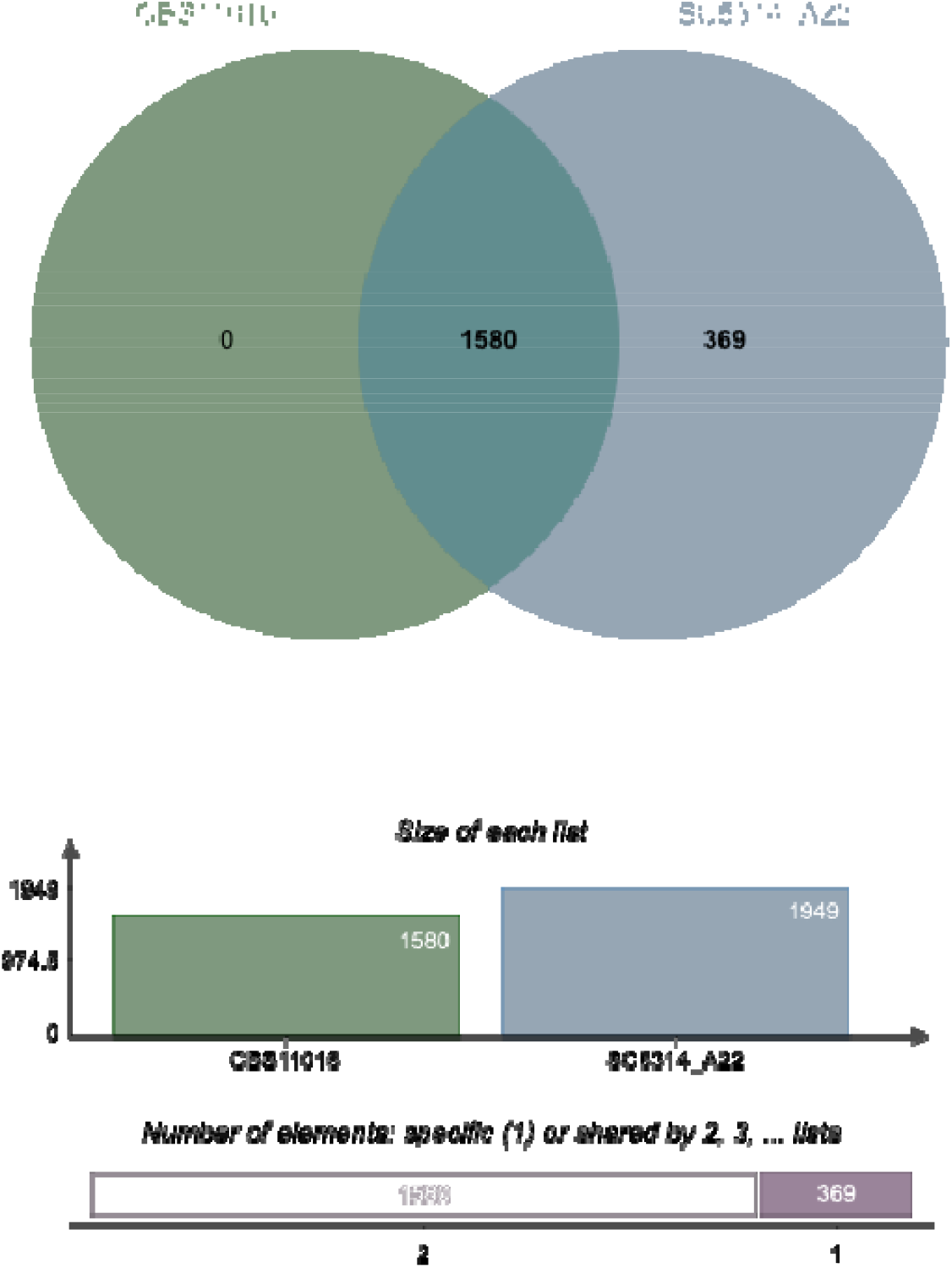
Venn diagram representing paralogous and orthologous groups between *C. albicans* and *C. africana* generated using Orthovenn2.

Comparative analysis showed a total of 331 structural variants (Figure 6) were found in the genome of *C. africana*.

**Figure 6.**
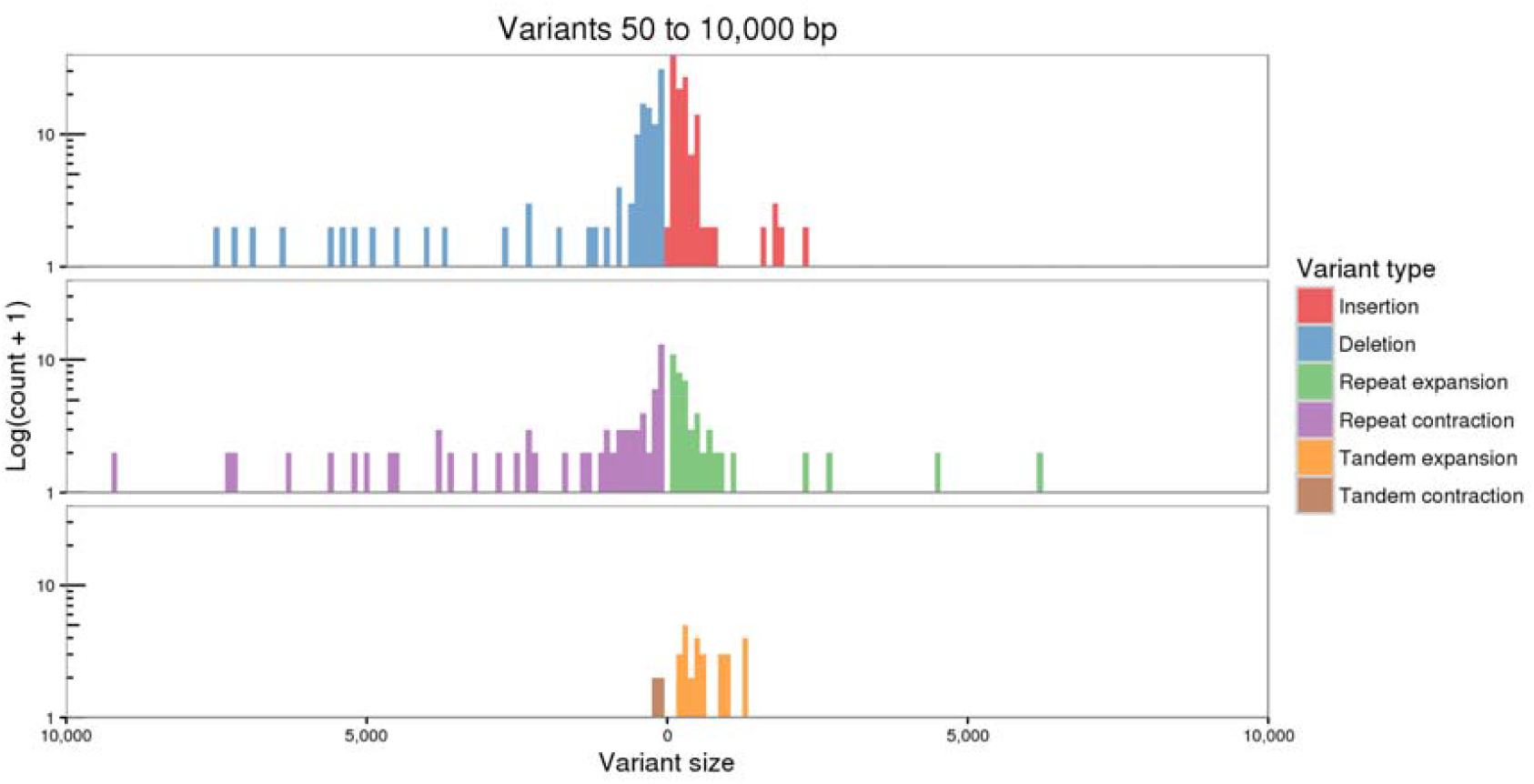
Structural variations between *C. africana* and *C. albicans* resulted from assemblytics[30].

## Discussion

This study is the first to study comparatively the genomes of *Candida africana* and *Candida albicans*. This is particularly interesting because the two species are similar but have different ecological niches and pathogenicity. In our analysis, we found that *C. africana* genome was comparable to the *C. albicans* genome both in terms of genome size and predicted coding genes (5934 coding genes in *C. africana* vs6221 coding genes in *C. albicans*). Orthovenn2[32] showed the presence of orthologous protein clusters across the genomes of *C. africana* and *C*. albicans. The identification of orthologous genes and ascertaining the degree of similarity between them are two important steps in comparative genomics studies to understand the evolution of genes and genomes[34]

Our orthologs analysis showed a total of 5557 orthogroups shared between *C. albicans* and *C. africana*, Interestingly, we found orthogroups exclusive to *C. africana*, containing a total of 12 genes coding for *C. africana*, whereas the orthogroups exclusive of *C. albicans* comprise a total of 35 genes whose product encoded.

### Evidence of Inversions in the genome of *Candida africana*

Comparing the genome of *Candida africana* and *Candida albicans* using syntenic dot plot. The presence of translocation of genes occurs in the genome. The genome of two organisms can differ due to evolutionary changes resulting in sequence mutations, chromosomal rearrangements, and gene family expansion or loss[35]. These gene rearrangements can be viewed using synteny Dot plot and Synteny mapping. Synteny identification is a filtering and organizing process of all local similarities between genome sequences into a coherent global picture[36]. Synteny blocks or collinearity are regions of chromosomes between genomes that share a common order of homologous genes derived from a common ancestor[37, 38]. Different tools were used to study evolutionary relationships between the genome of *C. africana* and *C. albicans*. SynMap[39], Synima[27] and progressive mauve[29].

Microsyntenic analysis of regions in the genomes of the two strains showed gene loss and gene gains in the genome of *C. africana*. The role of this gene loss or gain play in driving evolution in *Candida africana* leading to the reduced virulence is yet to be evaluated.

Microsyntenic analysis of regions highlighted in Figure 4. Each panel shows two genomic regions, *Candida africana* on top and *Candida albicans* on the bottom. A dashed line separates the top and the bottom strands of DNA in each panel. Coding sequences are shown in green, transcripts in blue, and full gene models in grey. Pink blocks identifying regions of sequence similarly identified by BlastZ and have semi-transparent wedges connecting them. If blocks are drawn on the top of the genomic region, the sequences are in the same orientation. If drawn on the bottom, they are in opposite orientations. Blocks with repeat sequences will be drawn on top of one another, and are visualized through the stacking of the transparent wedges.

The genomes of *C. africana* and *C. albicans* show a great level of synteny with the presence of translocations, deletions, and inversions. To compare regions in the dot plot that shows break and inversion, a GEvo[37] plot was performed on that region. Gevo plot reveals inversions occurring within the genome of *Candida africana*. Synonymous mutation(ks) values show that the genome of *C. africana* has not experienced recent duplication events.

### The presence of Structural variations exists between *C. africana* and *C. albicans*

Structural variations(SV)include insertions, duplications, deletions, inversions, and translocations that occur within a genome[40], Structural variations are aligned variations occurring between two aligned genomes with a size of above 50bp[41]. While present in a genome, SV can disrupt gene functions [42] In *Candida albicans*, single nucleotide polymorphisms And large scale variations can have effects in new clinical phenotypes[43], genomic variation is also well established for *C. albicans* [44].

### Conclusion

The genome of *C. africana* has lots of structural variations and the presence of gene losses and gains. These genetic variations possibly play a role in the reduced virulence potential observed in *C. africana*. This study highlights the presence of genetic variations within the genome of *C. africana* that might be responsible for the reduced virulence and ecological niche restriction in the genome of *Candida africana*. The study calls for further evaluation of these variations using appropriate models.

## Acknowledgment

We wish to thank Dr. Christophe d’Enfer et al., for permitting us to use the whole genome sequencing data of the C. africana CBS 11016 strain and for their valuable advice. We wish to appreciate Dr. Farrer Rhys, from the University of Exeter for his technical direction on the use of Synima and in the conception of this project. We also wish to thank Eric Lyons for support in comparative genome assembly using GEVo on CoGe platform. This work is supported by a grant to Nnadi, N.E by the Africa Centre of Excellence in phytomedicine research and development.

## DATA AVAILABILITY

Genome is available under the accession number JAENJO000000000 and bioproject number PRJNA432884

